# Haldane’s Probability of Mutant Survival is Not the Probability of Allele Establishment

**DOI:** 10.1101/704577

**Authors:** Ivan Krukov, A.P. Jason de Koning

**Author notes:** Corresponding authors: University of Calgary, Cumming School of Medicine, Department of Biochemistry and Molecular Biology. Email: (IK), (APJdK).

## Abstract

Haldane notably showed in 1927 that the probability of fixation for an advantageous allele is approximately 2*s*, for selective advantage *s*. This widely known result is variously interpreted as either the fixation probability or the establishment probability, where the latter is considered the likelihood that an allele will survive long enough to have effectively escaped loss by drift. While Haldane was concerned with escape from loss by drift in the same paper, in this short note we point out that: 1) Haldane’s ‘probability of survival’ is analogous to the probability of fixation in a Wright-Fisher model (as also shown by others); and 2) This result is unrelated to Haldane’s consideration of how common an allele must be to ‘probably spread through the species’. We speculate that Haldane’s survival probability may have become misunderstood over time due to a conflation of terminology about surviving drift and ‘ultimately surviving’ (*i.e.*, fixing). Indeed, we find that the probability of establishment remarkably appears to have been overlooked all these years, perhaps as a consequence of this misunderstanding. Using straightforward diffusion and Markov chain methods, we show that under Haldane’s assumptions, where establishment is defined by eventual fixation being more likely that extinction, the establishment probability is actually 4*s* when the fixation probability is 2*s*. Generalizing consideration to deleterious, neutral, and adaptive alleles in finite populations, if establishment is defined by the odds ratio between eventual fixation and extinction, *k*, the general establishment probability is (1 + *k*)/*k* times the fixation probability. It is therefore 4*s* when *k* = 1, or 3*s* when *k* = 2 for beneficial alleles in large populations. As *k* is made large, establishment becomes indistinguishable from fixation, and ceases to be a useful concept. As a result, we recommend establishment be generally defined as when the odds of ultimate fixation are greater than for extinction (*k* = 1, following Haldane), or when fixation is twice as likely as extinction (*k* = 2).

## Introduction

Established alleles are those that have persisted long enough in a population so that they are unlikely to be lost by chance, and can thus be said to have escaped loss by drift. The concept of establishment traces back at least to Fisher (1922, p. 419). Fisher argued that the fate of a new positively selected mutant is determined more by random drift when it is rare than if it persists long enough to spread through much of the population. Later, Haldane (1927) sought to “consider the course of events in a population where the new factor is present in such numbers as to be in no danger of extinction by mere bad luck”. In that work, Haldane used a branching process formulation to derive his famous probability of eventual fixation for a beneficial mutant with selection coefficient *s* (*P*_Fix_ ≈ 2*s*), which was termed the “probability of survival”. Perhaps because of context and the use of this term, which comes from the theory of branching processes, 2*s* is now commonly interpreted as both the fixation probability (*e.g.*, Kimura 1962; Otto and Whitlock 1997) *and* as the establishment probability (*i.e.*, the probability that an allele will reach a sufficiently high frequency to have effectively escaped loss by drift) (*e.g.*, Gerrish and Lenski 1998; Peck 1994; Messer and Petrov 2013). However, while fixation implies prior establishment, establishment should not necessarily imply fixation with complete certainty, or else it is at best a redundant concept. The probability of establishment should thus nearly always be greater than the probability of fixation, and the difference between the two should vary according to how exactly we define establishment.

A reading of Haldane’s paper (Haldane 1927) supports this view. In particular, after deriving the probability of fixation, Haldane went on to argue that a dominant allele achieving a population count of log(2)/2*s* will “probably” spread through the species. Although it was not stated explicitly, this is an approximation to the number of initial mutants that make the probability of fixation equal to 1/2, so that exceeding this threshold implies better than even odds that the allele will go on to become fixed (derivation below). Here we clarify that, unlike in the branching process formulation (Ewens 2004, p. 29), fixation and survival to establishment can be easily differentiated in a Wright-Fisher model. A Wright-Fisher perspective will thus clarify that Haldane’s 2*s* is unrelated to his establishment count, since the probability of fixation given the establishment count of log(2)/2*s* is 1/2 (not 2*s*). Under Haldane’s assumptions, we will see that the establishment probability of a beneficial allele is rather approximated by 4*s*, and is therefore off by a factor of 2 compared to the probability of fixation. This result applies to both haploid and diploid populations with zygote/heterozygote fitness of 1 + *s*. When heterozygote fitness is defined as (1 + *hs*) for *h* = 1/2, the probabilities of fixation and establishment are rather *s* and 2*s* respectively.

We begin by reviewing Haldane’s treatment and its assumptions. We next consider some related arguments by Gillespie (2004), before moving to a diffusion approach and eventually to a direct analysis of the discrete-time Wright-Fisher model (where we are not required to assume weak mutation, weak selection, or large population size). Moving to a full Markov chain treatment is important for validation, and because the cases where establishment are most of interest occur when population mutation rates may be very large and thus could violate assumptions that diffusion approximations usually require (de Koning and de Sanctis 2018). We focus on the general case where mutants may be deleterious, neutral, or advantageous and show that a diffusion approximation to the establishment probability has a pleasing simplicity when defined appropriately. We conclude that both the probability and rate of establishment (Messer and Petrov 2013) are different from what has been previously understood by as much as a factor of 2.

### Establishment count in a branching-process model

Haldane used a branching process formulation to consider the ultimate survival of a mutant allele. Assuming individuals leave a Poisson-distributed number of offspring in the next generation, and that the number of mutant offspring has mean 1 + *s*, the probability that the mutant population will eventually go extinct in the limit of *t* → ∞ can be determined by considering the probability that a newly arisen mutant in the current generation, *t*, will eventually go extinct, *P*_Ext_(*X*_*t*_ = 1) (where *X*_*t*_ indicates the number of mutants present in generation *t*). Noting that *P*_Ext_(*X*_*t*_ = 1) is equivalent to 1 − *P*_Fix_(*X*_*t*_ = 1), this can be expressed by writing the probability that an allele will leave *i* mutant offspring in the next generation times the probability that every one of the *i* mutants will eventually go extinct, integrated over all *i*:

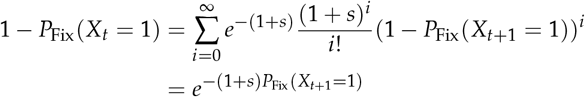

By noting that the probability of fixation of a newly arisen mutant should be the same in each generation under constant population size and selective effect (Otto and Whitlock 1997), we can set *P*_Fix_(*X*_*t*_ = 1) and *P*_Fix_(*X*_*t*+1_ = 1) to *P*_Fix_(*X*_0_ = 1) and solve for *s* in terms of *P*_Fix_. This yields

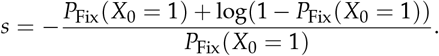

Suppressing the dependence on *X*_*t*_ for convenience, we take the Taylor series expansion of the solution for *s* around *P*_Fix_ = 0,

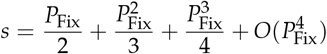

and obtain Haldane’s famous result for the probability of fixation of a beneficial allele when using the leading term:

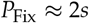

As noted above, Haldane next derived a minimum allele count, *c*^∗^ = log(2)/2*s*, that if exceeded would ensure that the allele will probably spread through the species (*i.e.*, establish). To see how he obtained this result, we begin with an approximation to the probability that *c* mutants will eventually go extinct based on the above result, (1 − 2*s*)^*c*^ (Haldane 1927; Otto and Whitlock 1997). This approximation works well when *s* is small and positive, and indeed it corresponds to Kimura’s extinction probability from diffusion theory for a starting frequency of *c*/(2*N*) in the limit of infinite population size (and to a first order approximation when *s* is small; not shown). To find the minimum number of starting copies such that fixation and extinction are equally likely, *c*^∗^, we use Haldane’s expression for the extinction probability and set this to 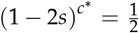, so that *c*^∗^ represents the count at which fixation becomes more likely than extinction when it is exceeded. Solving for *c*^∗^ yields

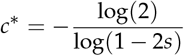

Taking the first term in a series expansion of this result around *s* = 0 then gives

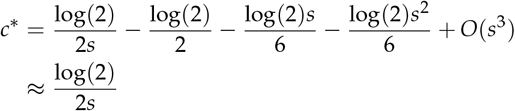

recovering Haldane’s result. We will refer to this quantity as Haldane’s establishment count. Throughout this paper, we refer to the minimum population frequency required to become established as the *establishment frequency* (*f*^∗^), and the corresponding allele count as the *establishment count* (*c*^∗^). Based on the above definitions, a population that achieves exactly Haldane’s establishment count will have a fixation probability of 1/2 (not 2*s*, which is the fixation probability assuming we started with a single mutant copy).

### Establishment frequency in a Wright-Fisher diffusion (*s* > 0, *N* → ∞)

Kimura (1962, 1964) considered the fixation probability in a diffusion approximation to a Wright-Fisher model including selection and drift in a series of celebrated papers. Given an initial mutant allele frequency *X*_0_ = *f*_0_, a population size of *N* diploid reproducing individuals, selection coefficient *s* and heterozygote fitness 1 + *s*, Kimura’s probability of fixation is given by:

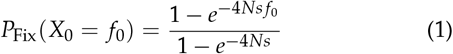

For a single starting copy (*f*_0_ = 1/(2*N*)), this yields

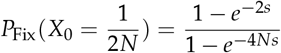

As Kimura (1962, eq. 11) noted, in the limit of infinite population size, this equation agrees with Haldane’s result to a first-order approximation:

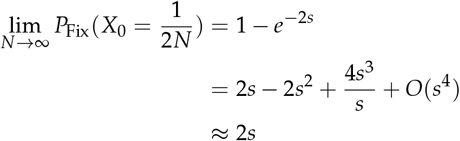

for *s* > 0 (where the series expansion was again taken around *s* = 0). This result supports the standard interpretation that Haldane’s 2*s* represents the fixation (not establishment) probability.

Using equation 1 as a starting point, Gillespie (2004, sec. 3.9, eq. 3.25) later derived an expression for the establishment frequency, *f* ^∗^, that makes the probability of fixation close to 1 (within a prescribed margin of error, *ε* ≈ 0, so that 1 − *ε* ≈ 1). Gillespie first assumed that 2*Ns* is large enough that the denominator of equation 1 approaches 1 and can be ignored. If we define heterozygote fitness as 1 + *s* and homozygote fitness as 1 + 2*s* (rather than the 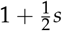 and 1 + *s* that Gillespie used), we obtain

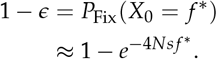

Solving for *f*^∗^, we then obtain

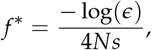

which is consistent with Gillespie’s reported result given our redefinition of heterozygote fitness. By 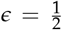 setting so that exceeding the establishment frequency makes the probability of fixation greater than the probability of extinction, this yields an establishment frequency that is equivalent to Haldane’s establishment count divided by 2*N*. Note that in both Haldane and Gillespie’s derivations, *s* was assumed positive and *N* was effectively assumed large so that the results do not necessarily apply to deleterious or neutral alleles, and do not account for finite population size effects.

We will now generalize the concept of allele establishment by combining the approaches of Haldane, Gillespie, and Kimura, and by relaxing assumptions about *s* and *N*. As we show, both the establishment frequency and probability have simple closed form approximations in the diffusion framework, even when the definition of establishment is varied according to the relative odds of eventual fixation to extinction.

## Results

Diffusion approximations to the establishment frequency and probability are now developed. These will be contrasted with numerical results obtained by independent analyses of discrete Wright-Fisher models, which are based only on the definitions of establishment and the model itself. The method used to efficiently analyze the discrete models is described in the Appendix.

### Establishment frequency in a Wright-Fisher diffusion (general case)

As above, we define the establishment frequency 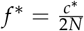 for a given establishment count, *c*^∗^. For generality, we define establishment in terms of the odds ratio *k*, such that eventual fixation is *k* times more likely than extinction. Definitions based on quantities other than the odds ratio are possible, however, as we will see, the odds ratio produces a convenient simplification and allows existing arguments as special cases (*e.g.*, *k* = 1 for Haldane, *k* = (1 − *ε*)/*ε* for Gillespie). Thus, the desired probability of fixation, *P*_Fix_(*X*_0_ = *f*^∗^), can be written as:

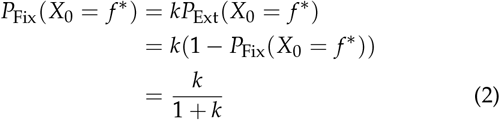

Following the approach used by Gillespie (2004), but without applying his approximations, equation 1 can be used to directly solve for the initial allele frequency *X*_0_ = *f*^∗^ that satisfies equation 2:

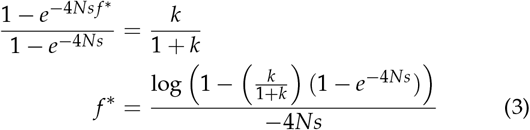

Immediately, we can confirm the intuitive result that when fixation and extinction are equally likely (*k* = 1), the establishment frequency (equation 3) of a neutral variant is 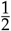:

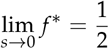

As a further check, multiplying the general solution in equation 3 by 2*N* and taking the limit as *N* → ∞ gives:

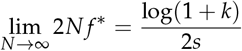

for *s* > 0. As expected, this result with *k* = 1 is equivalent to Haldane’s establishment count when *s* > 0.

### Establishment probability in a Wright-Fisher diffusion

Establishment probability can be defined as the probability of reaching the establishment frequency (or count) before going to extinction. Establishment may therefore be considered in a two absorbing-state Wright-Fisher model, with absorbing states defined at *X*_*t*_ ∈ {0, *f*^∗^} (in the continuous state-space). An expression for the establishment probability can then be found using diffusion theory in a manner analogous to how Kimura derived the fixation probability, but where the fixation absorbing boundary is moved to the establishment frequency and its meaning redefined.

Following Kimura (Kimura 1962), let *u*(*f*_0_, *t*) be the probability that the mutant allele will absorb at the upper absorbing boundary during time interval *t*, given initial frequency *f*_0_. We are interested in a solution to lim_*t* → ∞_ *u*(*f*_0_, *t*) = *u*(*f*_0_), with the boundary conditions *u*(0) = 0 and *u*(*f*^∗^) = 1. We solve for *u*(*f*_0_) by setting up the appropriate Kolmogorov backward equation, ignoring higher-order terms, and solving the ordinary differential equation:

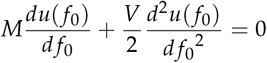

Under a diploid Wright-Fisher model accounting for drift and selection, the effective per generation mean and variance of the diffusion are *M* = *sp*(1 − *p*) and *V* = *p*(1/*p*)/(2*N*), respectively (when heterozygote fitness is 1 + *s*)). With the appropriate boundary conditions defined above, the solution is then

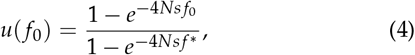

which has an obvious similarity to the fixation probability in equation 1. Substituting in the establishment frequency from equation 3, we obtain the simple result:

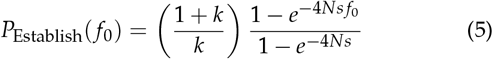

or simply (1 + *k*)/*k* · *P*_Fix_(*f*_0_). While *f*_0_ will usually be assumed to be 1/(2*N*), it need not be so and could reasonably take any value between 1/(2*N*) and *f*^∗^ − 1/(2*N*). This result also holds for dominant and recessive variants (not shown for brevity).

### Establishment and fixation probabilities differ according to the definition of establishment

Establishment and fixation probabilities differ according to how certain we wish to be that establishment implies likely fixation (Figure 1). On the left, establishment is defined as exceeding the frequency at which eventual fixation and extinction are equally likely (*k* = 1). As explained in the introduction, this roughly corresponds to the assumptions of Haldane (although he considered only the case of *s* > 0 in a branching-process model). With *k* = 1, the probability of establishment is twice the probability of fixation, while with *k* = 2 (where fixation is twice as likely as extinction), the establishment probability is 1.5× larger than that for fixation. On the right of Figure 1, establishment is defined such that fixation is extremely likely (100 times more likely than extinction, *k* = 100). This corresponds to Gillespie’s definition of establishment with *ε* set to ~0.01 (1/101 to be precise). Comparison of a direct analysis of the discrete Wright-Fisher model with the diffusion results of the previous sections confirms the results in equation 5. Since the direct analysis of the discrete model is agnostic to details of the model itself, results for several values of the population mutation rate parameter, *θ*, were computed where both forward and backward recurrent mutation were allowed. The close correspondence of these results with the diffusion results (which assumed no new mutations) suggests that the relationship between *P*_Fix_ and *P*_Establish_ is robust as defined in equation 5.

**Figure 1.**
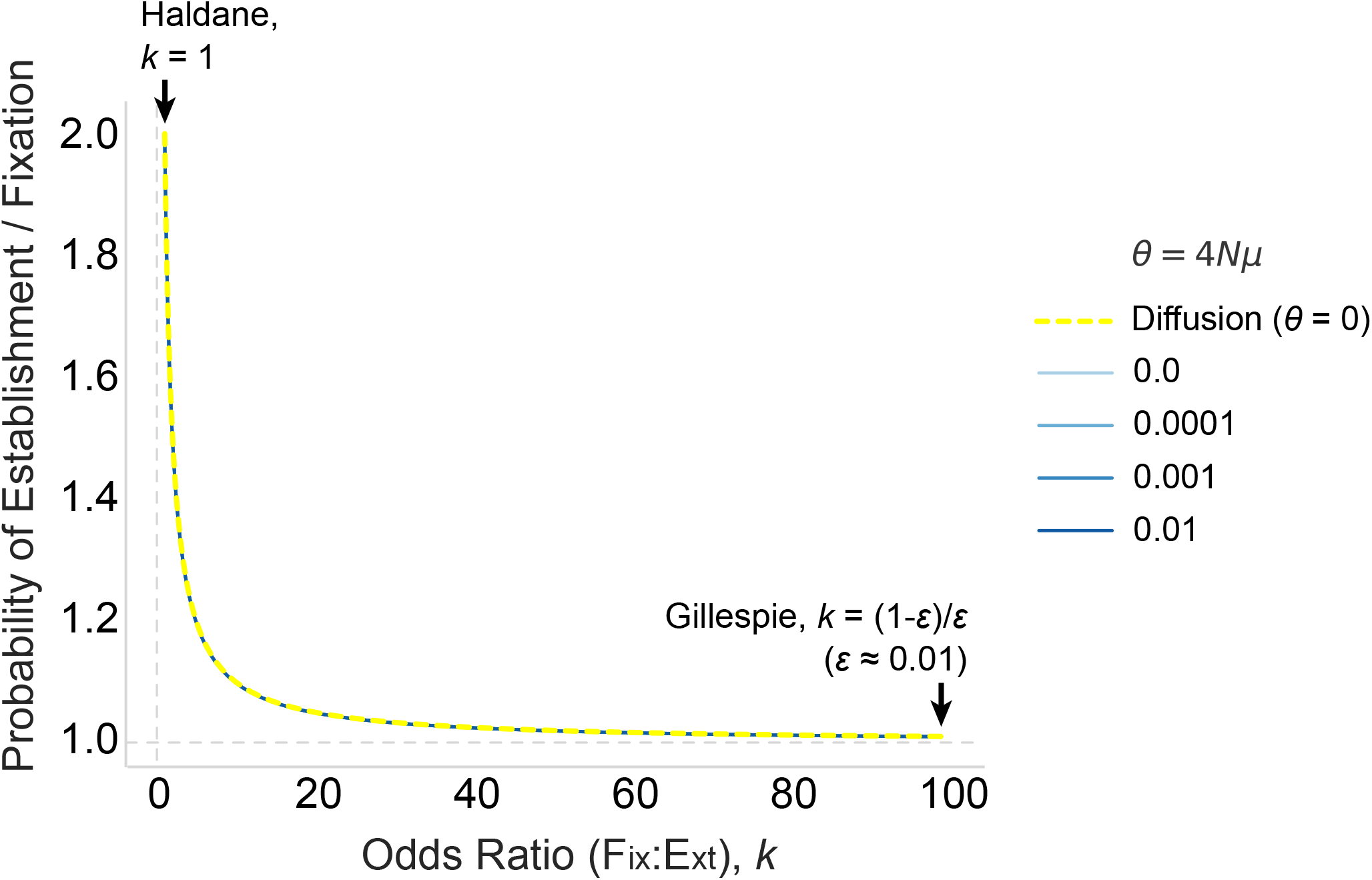
Establishment probability compared to fixation probability for a range of fixation-to-extinction odds ratios (*k*). The dashed yellow line shows results of calculations made assuming *p* = 1/(2*N*) and using the diffusion method. Solid lines show results based on direct computation of the establishment count and its corresponding probability using the discrete-time Wright-Fisher model. Note, the methods agree and numerically correspond with (1 + *k*)/*k* as implied by equation 5. Accordingly, the results are invariant to *N* and *s*; these results were computed for *s* = 0 and *N* = 10, 000. Values of *k* = 1 and *k* = (1 − *ε*)/*ε* capture the assumptions of Haldane and Gillespie, respectively. At Haldane’s more liberal definition of establishment, the probability of establishment is twice the probability of fixation, whereas for Gillespie’s definition, *P*_Establish_ → *P*_Fix_ approximatlely as *ε* → 0).

In Figure 2, establishment frequency and probability are contrasted under Haldane’s definition of establishment (*k* = 1), using both the general diffusion solutions presented above, and the direct analysis of the discrete Wright-Fisher model. Establishment frequency is large and close to one for deleterious alleles, illustrating that strongly deleterious alleles are only expected to establish in an equilibrium population when they start at frequencies that are already very high. Contrariwise, beneficial alleles have comparatively low establishment frequencies, which decrease as the strength of positive selection is increased. For neutral mutations, the establishment frequency is 0.5, and the shape of the curve is symmetric with respect to the neutral establishment frequency. Only minor variations are observed when bidirectional recurrent mutation is introduced (where faster mutation generally exaggerates the effects of selection).

**Figure 2.**
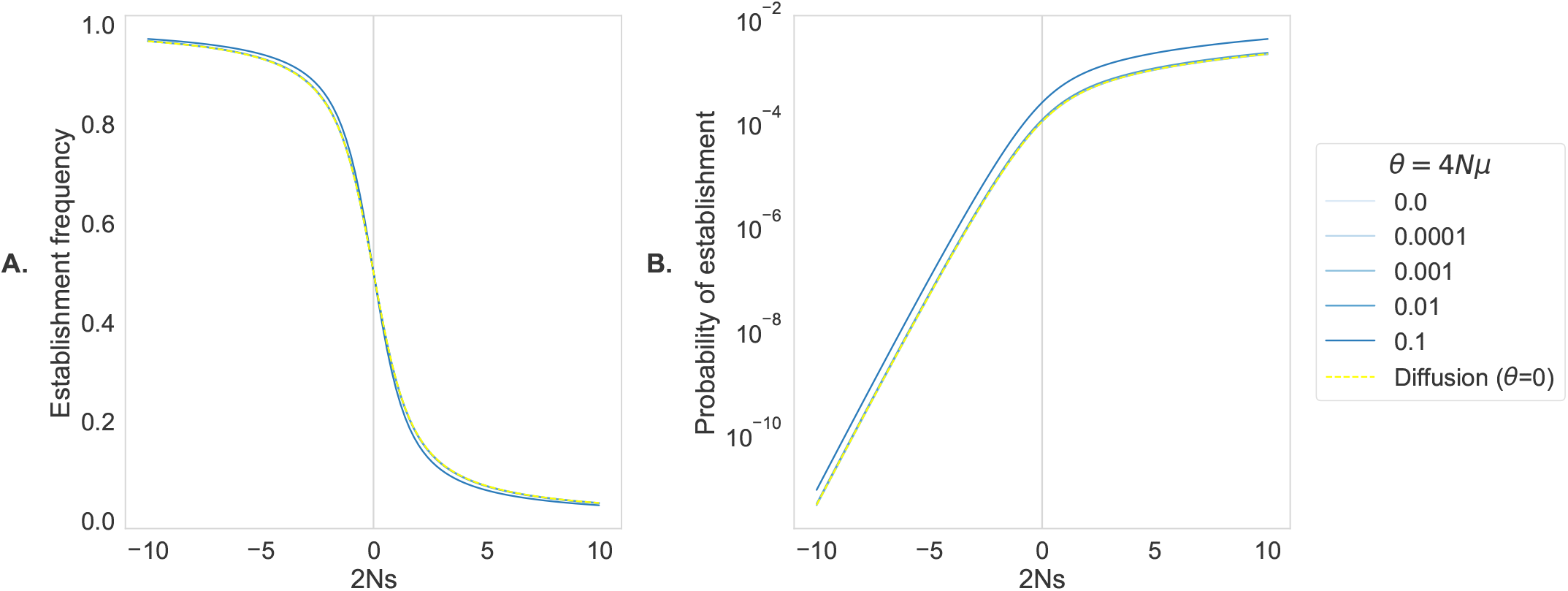
Comparison of establishment frequencies and probabilities for *k* = 1. As before, dashed yellow lines show analytic diffusion solutions, solid blue lines show discrete solutions. The diffusion approximation agrees with the discrete solution in this parameter range. (A) Establishment frequency as a function of selection. At neutrality, establishment frequency is 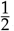, increasing for negatively selected, and decreasing for positively selected alleles. (B) Probability of establishment as a function of selection (Yaxis on log scale). Establishment probability of a neutral variant is 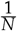, increasing for positively selected, and sharply decreasing for negatively selected alleles. Population size *N* = 10, 000, heterozygote fitness (1 + *s*)

In the discrete framework, we can also easily find the expected time to establishment (for those variants that establish; Figure 3A), the expected time of segregation after establishment (Figure 3B), and the expected time to fixation after establishment (for those variants that go on to be fixed; Figure 3C). These quantities could also be found using diffusion theory for particular model parameterizations, which we do not pursue here for brevity. As expected, the establishment time for advantageous alleles is shortest (Figure 3A). However, it is interesting to note that mildly deleterious alleles take a significantly longer time to establish than do either neutral or strongly deleterious alleles. Similar effects have been observed in computations of expected allele age and times to absorption, and have been attributed to ‘stochastic slowdowns’ under weak selection in the presence of dominance (Mafessoni and Lachmann 2015; de Sanctis *et al.* 2017) and mutation (de Sanctis *et al.* 2017). Notably, we see these effects here even when the mutation rate is zero. Similarly, we observe that once established, weakly adaptive alleles take longer to go to fixation than do neutral alleles (Figure 3C). Contrariwise, the average time to absorption (at either boundary) after establishment is symmetric with respect to selection about *s* = 0 (Figure 3B).

**Figure 3.**
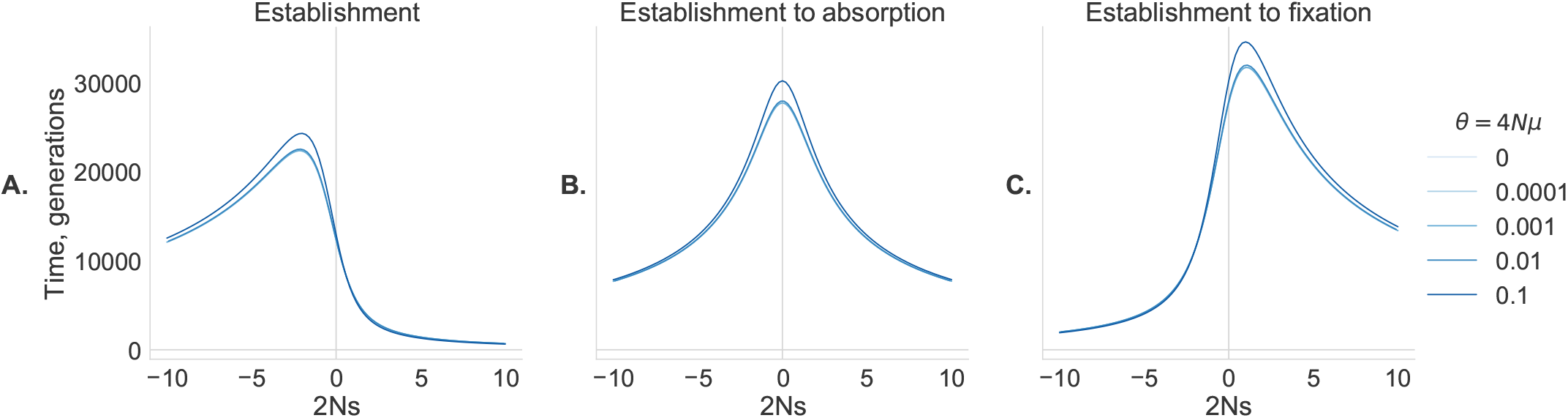
Expected times to establishment and absorption in the discrete Wright-Fisher model with establishment defined by *k* = 1. (A) Expected time to establishment (from *f*_0_ = 1/(2*N*)); (B) Expected time to absorption, post-establishment (at either absorbing boundary); and (C) Expected time to fixation, post-establishment. Solid blue lines correspond to different values of the bidirectional mutation rate. Population size *N* = 10, 000, heterozygote fitness (1 + *s*).

## Discussion

We have shown that establishment (survival until escaping loss by drift) and fixation (taking over the population) are easily differentiated in a Wright-Fisher model, and that clearly defining the establishment frequency and probability clarifies the meaning of Haldane’s important result (*P*_Fix_ ≈ 2*s*, *s* > 0). Despite that Haldane’s fixation probability is widely interpreted as if it were the establishment probability, we have argued that the probability of reaching the establishment frequency does not appear to have been addressed until now (except in the trivial case where *k* is large and establishment therefore implies certain fixation). Accordingly, we do not believe Haldane intended his probability of fixation to be interpreted as the probability of reaching the establishment count. Indeed, it is plausible that his result has been misinterpreted owing to the use of terminology from the theory of branching processes that may be ambiguous when taken out of context. For example, Haldane called 2*s* the ‘probability of survival’, alluding to the so-called ultimate survival probability in branching process theory. While this sounds very much like survival to escape of loss by drift, which he also discussed, it really means fixation since it refers to the ultimate fate of an allele in the limit of infinite (or very long) time.

We recognize that there could be reasonable disagreement amongst geneticists about the definition of establishment in comparison to fixation. After all, historical usage of the term ‘establishment’ has been inconsistent, with some authors using it as a synonym for complete fixation (*e.g.*, Kimura and Ohta 1969) and others using it specifically to mean survival to escape from loss by drift (*e.g.*, Haldane 1927). Nevertheless, the modern usage is more consistent with the latter definition, which we have assumed throughout this work. Due to this ambiguous legacy, it would probably be helpful to avoid future colloquial uses of the term ‘establishment’, *sensu lato*, without explicitly defining what is intended.

Defining establishment with respect to the odds ratio of eventual fixation to extinction provides a flexible definition that generalizes previous approaches, and which allows the establishment probability to be simply computed as (1 + *k*)/*k* times *P*_Fix_. Under Haldane’s assumptions, this corresponds to an establishment probability of 4*s* (when *P*_Fix_ = 2*s*; *k* = 1), and 3*s* when establishment is defined by fixation being twice as likely as extinction (*k* = 2). When heterozygote fitness is 1 + *sh* and *h* = 1/2, the establishment and fixation probabilities are half of these values (not shown). For larger values of *k* (*e.g.*, *k* > 10), *P*_Establish_ is quite close to *P*_Fix_ and the concept itself becomes equivalent to fixation. We therefore suggest that *k* = 1 and *k* = 2 are reasonable choices for defining establishment in practice.

## Acknowledgments

We thank Bianca de Sanctis for helpful comments and discussion. This work was supported by a Discovery Grant to APJdK from the Natural Sciences and Engineering Research Council of Canada (NSERC), and by an Alberta Innovates Technology Futures (AITF) Doctoral Fellowship to IK. The authors gratefully acknowledge infrastructure support from the Canada Foundation for Innovation, Alberta Advanced Education, and the Alberta Children’s Hospital Research Institute.

## Appendix

### Establishment count in a Wright-Fisher Markov model (general case)

Here we derive establishment properties for arbitrary discrete-time Wright-Fisher models. Detailed discussion of efficient computational solutions to the discrete-time model can be found in Krukov *et al.* (2017); de Sanctis *et al.* (2017); de Koning and de Sanctis (2018).

Consider a discrete-time Wright-Fisher model with the transition probability matrix:

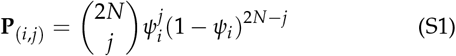

States 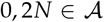 are absorbing states, corresponding to extinction and fixation, respectively. The rest of the states are transient, 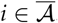. The transition probability matrix (equation S1) can be partitioned into transient-to-transient transition probabilities 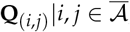, and transient-to-absorbing transition probabilities 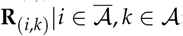:

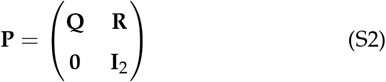

In this case, **R**= [**R**_Ext_, **R**_Fix_] has two columns, corresponding to extinction and fixation. The full model has 2*N* + 1 states, corresponding to allele counts from 0 to 2*N*, inclusive. The fundamental matrix of the Markov chain is:

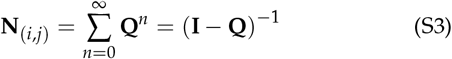

The entries **N**_(*i*,*j*)_ express the expected number of generations spent in each state *j* prior to absorption, conditional on starting in state *i*. The probability of absorbing in each absorbing state, conditional on starting in state *i*, is given by the matrix:

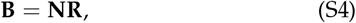

which has one column of probabilities for each of the absorbing states.

We can find the establishment count *c*^∗^ directly by scanning **B** for increasing values of *c*_0_ (the initial state), until we find the first entry of **B**’s second column such that 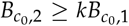. Note that this does not require solving for every row of **N**, since we can rearrange

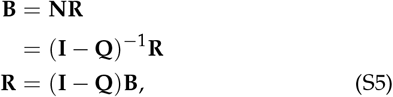

which can be easily computed using an LU decomposition of the matrix **I** − **Q**.

### Probability of establishment in a Wright-Fisher Markov model (general case)

Given the establishment count *c*^∗^, we can define a new Markov model with *c*^∗^ as the upper absorbing boundary, yielding a model with *c*^∗^ − 1 × *c*^∗^ − 1 transient-to-transient state transitions. This reduced model is constructed by truncation of **Q**, and by setting the second column of *R* to 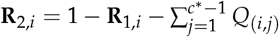.

With the full transition matrix **P**′ thus defined, where we call the corresponding transient-to-transient submatrix **Q**′, and the transient-to-absorbing submatrix **R**′, we can find the properties of interest by using **N**′ = (**I** − **Q**′), as in equation S3. We can then use **N**′ to derive properties such as probabilities and expected times using standard definitions (Krukov *et al.* 2017; de Sanctis *et al.* 2017). The matrix **Q**′ (just as matrix **Q**) can be based on any parameterization of the underlying model, including with arbitrary mutation, dominance, and selection.

To integrate quantities of interest over the likely distribution of starting states **c**_**0**_, which can become important when the population mutation rate is not small, we integrate over each state according to the probability of mutation creating 1, 2, 3, … copies in a single generation, starting from zero mutant copies (*i.e.*, *P*_0,1_, *P*_0,2_, …). As in de Sanctis *et al.* (2017) and de Koning and de Sanctis (2018), the summation can be truncated at *x* terms for *x* when *P*_0,*x*_ falls below some small threshold (*e.g.*, 10^−7^).

